# Transcriptome analysis of circulating microRNAs associated with Chagas disease susceptibility and chronic Chagas cardiomyopathy severity

**DOI:** 10.1101/2025.08.27.672539

**Authors:** Eric Henrique Roma, Fabielle Marques-Santos, Alinne Rangel dos Santos Renzetti, Luciana Fernandes Portela, Filipe Pereira da Costa, Alejandro Marcel Hasslocher-Moreno, Marcelo Teixeira de Holanda, Luiz Henrique Conde Sangenis, Andréa Rodrigues da Costa, Joseli Lannes-Vieira, Tania Cremonini de Araujo-Jorge, Roberto Rodrigues Ferreira, Roberto Magalhães Saraiva, Maria da Glória Bonecini-Almeida

## Abstract

**Background:** Chagas disease (ChD), caused by infection with the protozoan parasite *Trypanosoma cruzi*, is a major public health concern in Latin America. Understanding the molecular mechanisms driving disease progression and identifying biomarkers are crucial.

**Objective:** We investigated the association of circulating microRNAs (miRNAs) with ChD susceptibility and chronic Chagas cardiomyopathy (CCC) progression.

**Methods:** A multicentric prospective observational study was conducted with 150 ChD patients (46 indeterminate form, 104 CCC staged A-D) and 42 non-ChD controls from ChD endemic areas from Rio de Janeiro, and Posse in Brazil. Sequencing of circulating miRNAs was performed, and differential expression analyses were conducted. The differentiated miRNAs were submitted to functional analyses related to immune response and cardiovascular signaling pathways.

**Results:** We identified 40 differentially expressed miRNAs between ChD patients and non-ChD controls, highlighting miR-199b-5p, miR-153-3p, miR-143-3p, and miR-223-3p upregulated, and miR-150-3p, miR-4508, miR-486-5p, and miR-3960, downregulated in ChD patients. Moreover, functional analysis using Ingenuity Pathway Analysis software revealed that these miRNAs were involved in immune response and cardiovascular signaling pathways. Several miRNAs (miR-6734-5p, miR-1285-5p, miR-10527-5p, miR-31-5p, miR-5187-5p, miR-6515-5p) tended to present a higher expression in patients with severe compared to those with mild CCC, while miR-30c-2-3p presented lower expression in these groups.

**Conclusion:** This study provides evidence for the dysregulation of specific miRNAs in ChD, highlighting their potential role in disease pathophysiology and possible use as biomarkers of ChD susceptibility and CCC severity.

## Background

Chagas disease (ChD), also known as American trypanosomiasis, is a neglected tropical disease caused by infection with the protozoan parasite *Trypanosoma cruzi*. This disease represents a significant public health problem in many countries of Latin America, mainly in areas with poor living conditions, social vulnerability, and close interactions with wild or domestic mammals that can harbor the parasite. ChD is transmitted to humans through the feces of triatomine bugs, also known as “kissing bugs”, which feed on human blood and deposit the parasites in their excreta, or transmission through oral, vertical, blood transfusion and organ transplant [1].

Around 7 million people may be infected with *T. cruzi*, most of them living in Latin America. Further, migration has brought ChD patients to North America, Europe and Asian countries [2]. It is estimated that 1.9-4.5 million people in Brazil are infected [3], being one of the most important endemic countries related to this disease.

ChD is characterized by a wide spectrum of clinical manifestations, ranging from an “indeterminate” form to severe cardiac and digestive complications. Approximately 30-40% of infected individuals will develop the chronic Chagas cardiomyopathy (CCC) form of the disease over the course of life, characterized by a myriad of manifestations that include, among the most important, supraventricular and ventricular arrhythmias, as well as atrioventricular blocks, all associated with sudden cardiac death; thromboembolic phenomena including stroke; and reduced left ventricular ejection fraction (LVEF) leading to heart failure [4,5]. Despite the well-known clinical consequences of ChD, the underlying molecular mechanisms that drive disease progression and the development of CCC are not fully understood. Moreover, biomarkers of disease progression are a demand to improve patient’s management.

MicroRNAs (miRNAs) are small, non-coding RNA molecules of 18-22 nucleotides that play a crucial role in the regulation of gene expression. Also, the expression of miRNAs might be regulated in a tissue-specific manner, such as cardiac tissue. Indeed, some miRNAs are associated with a potential role in cardio-protection, modulating cardiac remodeling, as well as inflammatory and fibrotic pathways, as miR-450a that was associated with cardiac protection in an ischemia-reperfusion injury model of isolated rat cardiomyocytes and in human AC16 cells [6]. Also, some miRNAs might be involved in cardiac damage, as increased expression of miR-208a has been linked to cardiac hypertrophy [7] and downregulation of miR-133 has been associated with the progression of cardiac fibrosis [8].

miRNAs have been implicated in the pathogenesis of various diseases, including cancer [9,10] and infectious diseases, such as malaria [11], COVID-19 [12], tuberculosis [13], and leishmaniasis [14]. In the light of our knowledge, few studies have explored the role of selected miRNAs in the different clinical presentations of ChD, including the indeterminate form and the different CCC stages [15–18]. However, only one study characterized circulating miRNAs from ChD patients using small RNA sequencing approach at the moment [18].

In the present study, we performed miRNA sequencing assay from blood samples of ChD patients, including those with the indeterminate form and with mild, moderate and severe CCC stages, in order to assess the association between miRNAs and ChD susceptibility and CCC severity. The results of this study may contribute to the identification of potential biomarkers for the early detection and risk stratification of ChD progression.

## Methods

### Patient cohort

This multicentric prospective observational study included patients diagnosed with chronic ChD who were referred to the outpatient clinic of the Evandro Chagas National Institute of Infectious Diseases (INI), Oswaldo Cruz Foundation, in Rio de Janeiro, Brazil. All patients at cohort inclusion were tested using an enzyme-linked immunosorbent assay (ELISA), and immunochromatography or chemiluminescence. Forty-two non-ChD controls (non-ChD) were enrolled with both negative confirmatory serological tests. These non-ChD controls were enrolled INI as well as in the city of Posse, Goiás State, in Brazil. Socio-demographic and clinical data were obtained from medical records and patient interviews. Adult participants of both genders who agreed to sign the informed consent form were enrolled from February 2022 to March 2023.

### Patients CCC classification

The degree of cardiac involvement was assessed using the Second Brazilian Consensus on Chagas Disease (2015) [19], which consider electrocardiogram (EKG) findings, echocardiogram (ECHO) results, and the presence of heart failure to categorize CCC severity as: stage A – altered EKG, normal ECHO without congestive heart failure (CHF); stage B1 – altered EKG and ECHO with LVEF ≥ 45%, without CHF; stage B2 – altered EKG and ECHO with LVEF ≤ 45%, without CHF; stage C – altered EKG and ECHO, with compensated heart failure; stage D – altered EKG and ECHO, with end-stage heart failure. To increase the statistical power of the analysis, we grouped the CCC patients as follows: subgroup A, patients in the stage A, corresponding to mild CCC; subgroup B1/B2, patients in the stages B1 or B2, corresponding to moderate CCC; subgroup C/D, patients in the stages C or D, corresponding to severe CCC.

Sample’s collection, miRNAs isolation, library preparation and sequencing.

Five mL of blood samples were collected from all participants, and serum was separated and stored at −80°C until miRNA isolation and sequencing analysis.

miRNAs from serum samples (n=192) were extracted using the miRNeasy Serum/Plasma Advanced kit (Qiagen, Hilden, Germany) following the manufacturer’s recommendations and stored at −80°C until use.

Five microliters of the extracted miRNA were used for library preparation for miRNA sequencing with the QIAseq miRNA Library kit (Qiagen) following the manufacturer’s recommendations. After library preparation, the quality and quantification of the libraries were assessed using the TapeStation system (Agilent, Santa Clara, USA) with the High Sensitivity DNA 1000 kit (Agilent). Subsequently, the prepared miRNA libraries were sequenced using 2 P3 NextSeq 1000/2000 50 cycles kits (Illumina, San Diego, USA) on the NextSeq 2000 platform (Illumina).

### Analysis of miRNA sequencing and function

The generated reads were trimmed and aligned with the human mirBase v22, and miRNA counts were generated using the CLC Genomics Workbench software version 25 (Qiagen). For differential expression analyses, the DESeq2 package [20] in the R environment was used.

The miRNAs functional analysis was predicted using Ingenuity Pathway Analysis (IPA) bioinformatics software (Qiagen). The miRNA function was filtered using the MicroRNA Target Filter tool, selecting pathways related to immune response and cardiovascular diseases.

### Statistical analysis

A descriptive analysis of clinical and demographic characteristics was performed. The Mann–Whitney U test evaluated quantitative variables, whereas the univariate Odds Ratio (OR) was used for categorical variables. The differential expression of miRNA between groups was adjusted for participants’ age and gender. The p-values of single comparison were corrected for multiple comparisons by False Discovery Rate (FDR). Principal component analysis was performed using the DEM from ChD versus non-ChD comparison. A p-value < 0.05 was considered significant. All analyses were performed using R software version 4.3.3 and Prism 10 (Graph Pad, San Diego, USA).

## Results

### Participantś characteristics

A total of 150 patients with ChD were divided into the indeterminate form (IND group, n=46) and the CCC group (n=104). The CCC group was further stratified into stage A (n=37, patients with mild CCC), stages B1/B2 (n=37, patients with moderate CCC), and stages C/D (n=30, patients with severe CCC). Additionally, 42 control individuals without ChD (non-ChD group) were recruited from endemic areas (Table 1).

**Table 1.**
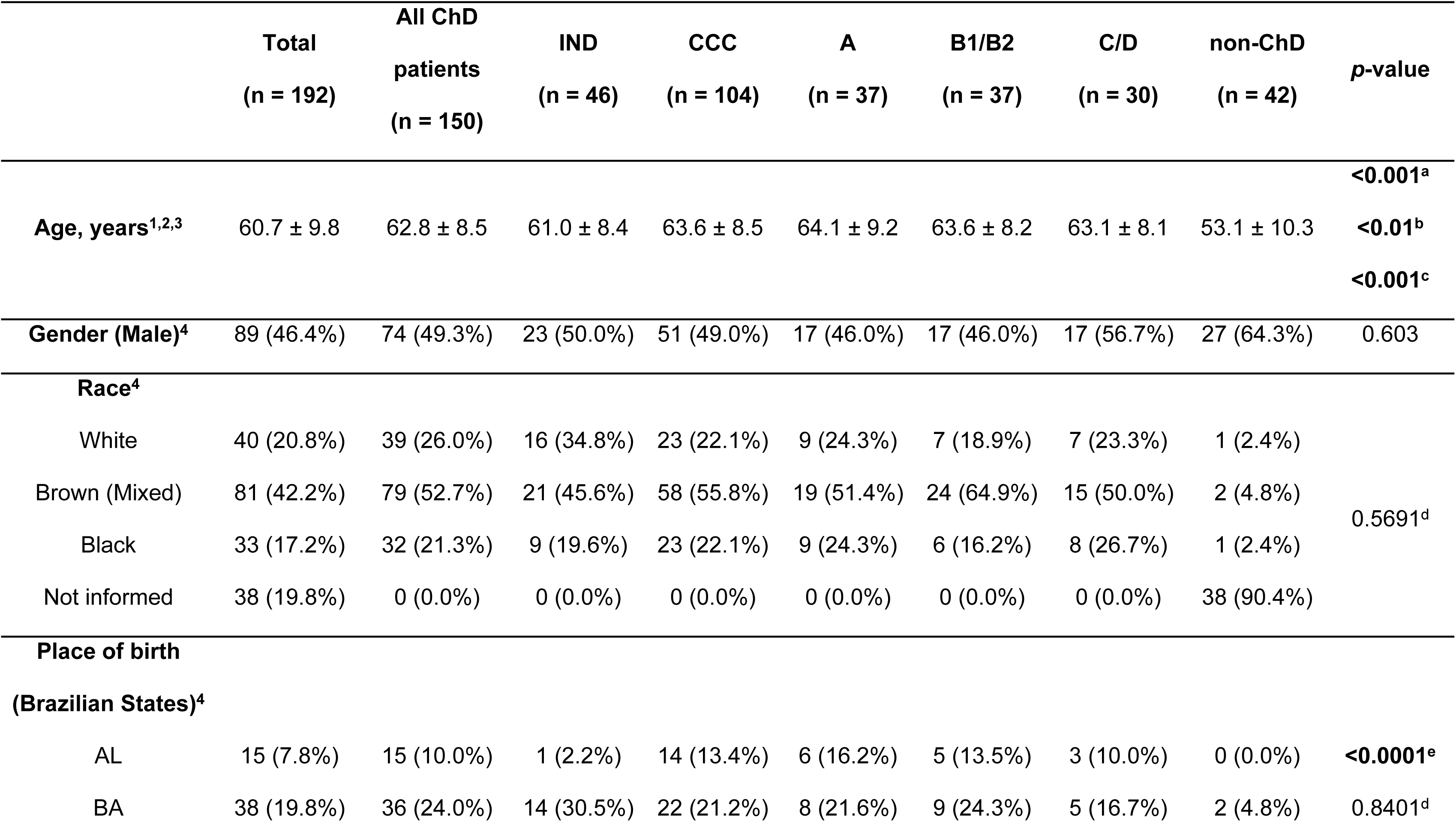
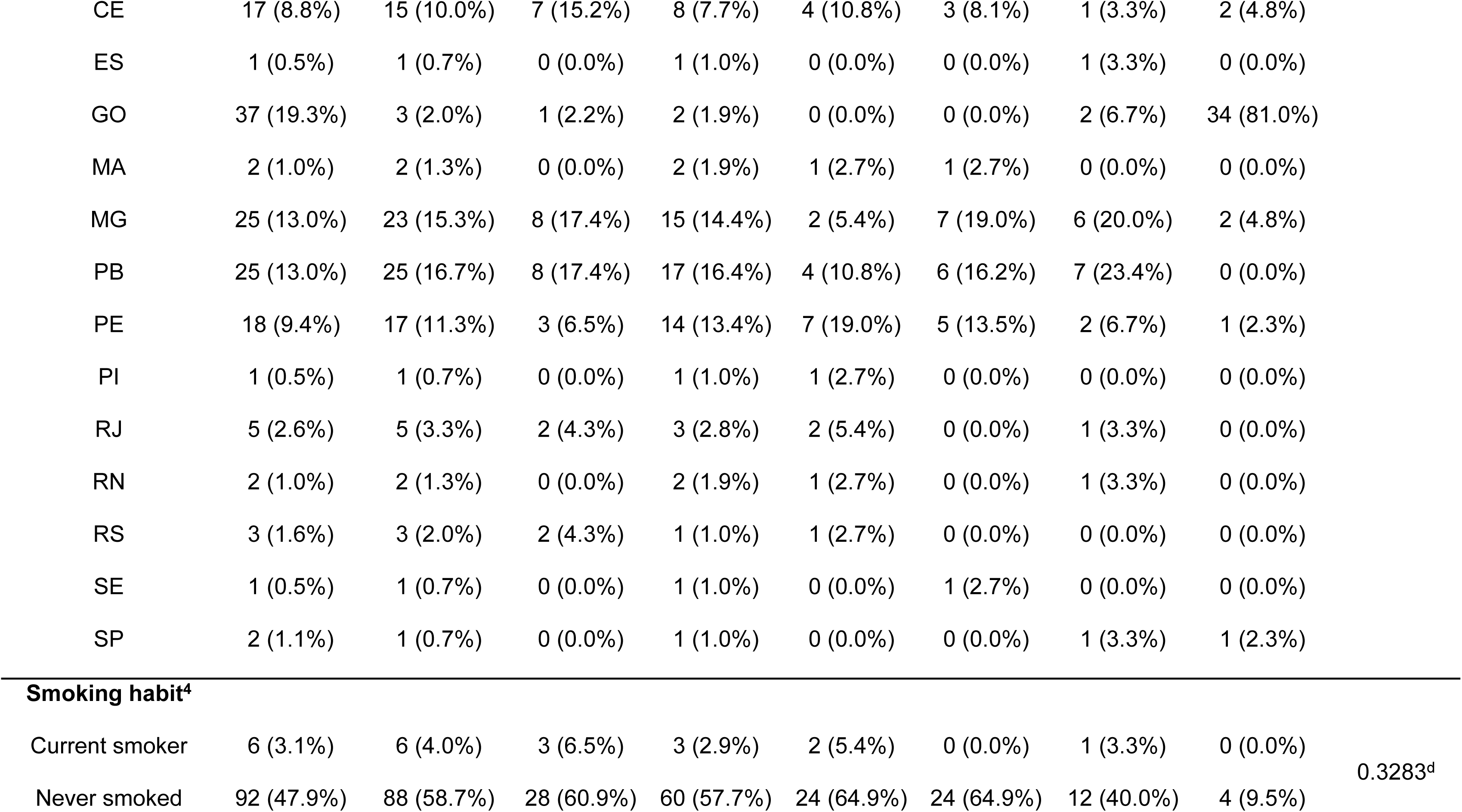
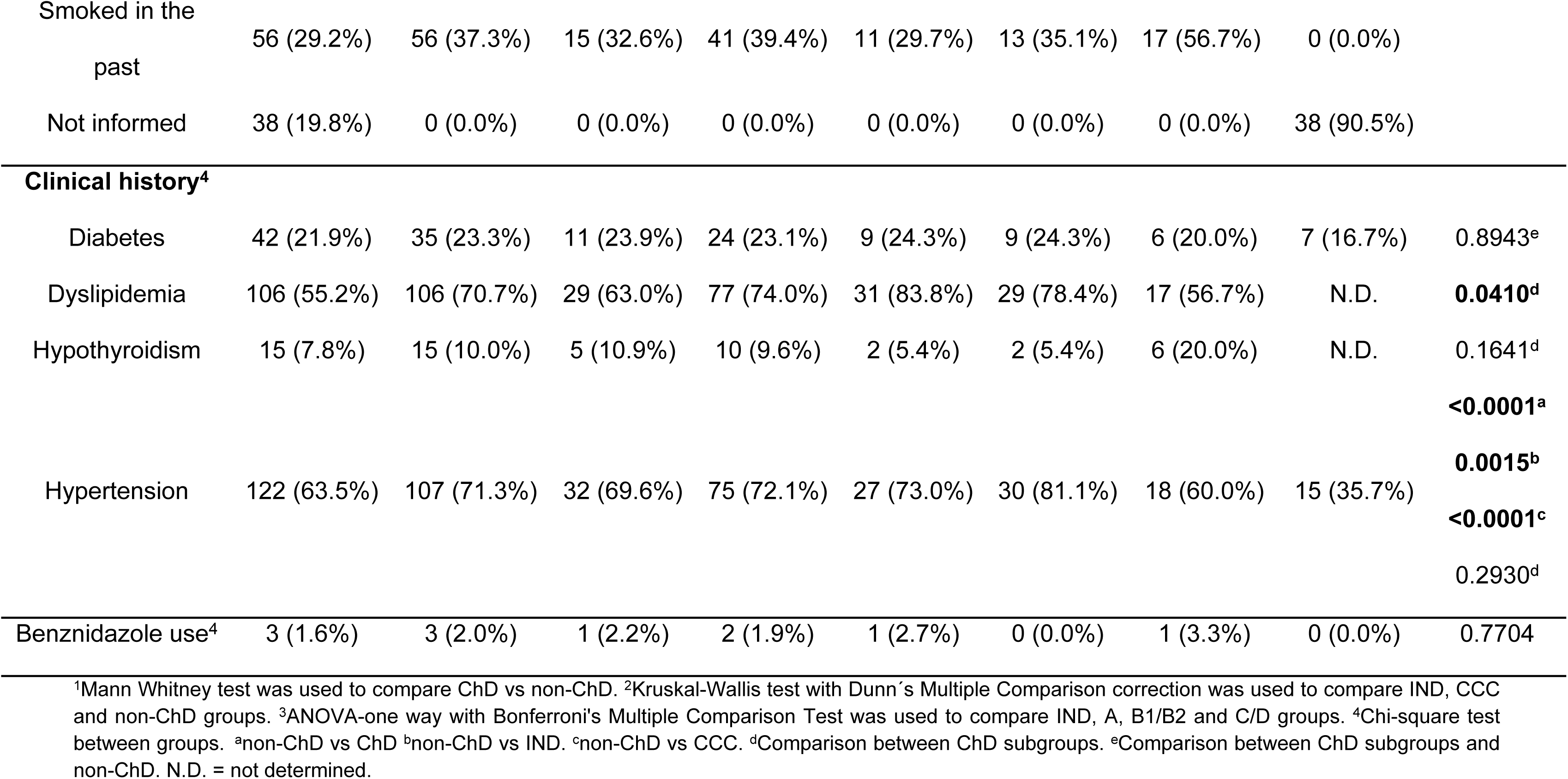
Patients’ characteristics.

ChD patients were significantly older than the non-ChD group (62.8 ± 8.5 versus 53.1 ± 10.3 years, respectively, p<0.001). This difference was sustained even splitting the ChD group in IND (61.0 ± 8.4 versus 53.1 ± 10.3, p<0.01) and CCC groups versus non-ChD (63.6 ± 8.5. versus 53.1 ± 10.3, p<0.001). There was no difference in the distribution of gender, race, and place of birth across the studied ChD groups. However, most of the non-ChD group originated from an endemic city of the state of Goiás (81%, p<0.0001), compared to ChD group and subgroups. Regarding the presence of comorbidities, only hypertension was more frequent in patients with IND form and CCC when compared to the non-ChD group (p<0.0015, and p<0.0001, respectively). In fact, the prevalence of hypertension in the ChD group (71.3%) was higher than in those of the non-ChD group (35.7%, p<0.0001).

### miRNA sequencing

Circulating miRNAs were sequenced with a mean of 5,119,952.47 ± 1,975,074.17 reads/sample and a mean of 86.20% ± 10.87% of the sample reads successfully trimmed. More than 98% of the sequencing showed a PHRED score >30, and Percent Passing Filter (%PF) of 83.72%. Eighty-six percent of the reads were successfully trimmed generating a mean of 14.87 ± 3.27 nucleotides read size, and 54.76% ± 26.99% of trimmed reads were annotated with record in miRbase v22.

### Differential expression analysis between ChD and control group

Initially, to identify miRNAs predictors of ChD susceptibility, we compared the miRNA expressions between ChD versus non-ChD groups. We found 144 differentially expressed miRNAs (DEMs) between both groups, considering a p-value of false discovery ratio (FDR) <0.05. After adjustment for age and gender and considering a log2 Fold Change <-1.5 and >1.5, and an FDR p<0.05, we observed 27 miRNAs upregulated, and 13 miRNAs downregulated indicating possible *T. cruzi* infection biomarkers (Fig. 1, supplemental Table 1). From these 40 DEMs, we performed cluster analysis of the selected 17 upregulated and 8 downregulated to identify possible different miRNA expression patterns among ChD subgroups. We observed a pronounced difference in miRNAs expression between ChD and non-ChD groups (Fig. 2), with distinguished pattern of DEMs in ChD patients compared to non-ChD groups as shown in principal component analysis (PCA) (Fig. 3A).

**Fig. 1:**
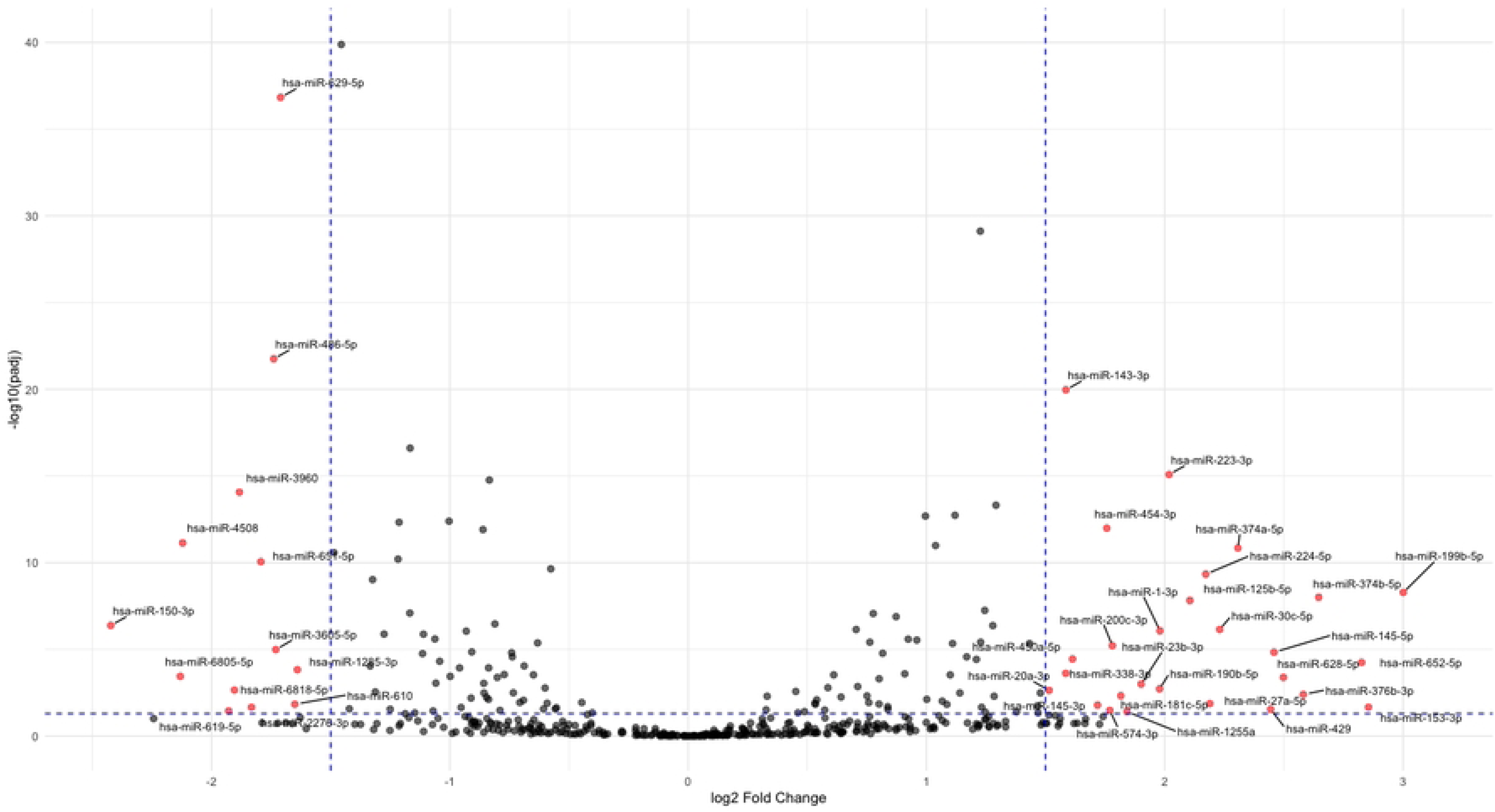
Differential expression of miRNAs (DEMs) in Chagas disease (ChD) patients compared to controls (non-ChD). Volcano plot showing differentially expressed miRNAs between ChD (n=150) and non-ChD (n=42) groups, adjusted for age and gender. The x-axis represents log2 fold change, and the y-axis represents the negative log10 of the *p*-value, corrected for multiple comparisons by False Discovery Rate (FDR). Red dots on the right side of the volcano plot indicate miRNAs significantly upregulated in ChD patients (log2 Fold Change > 1.5, FDR *p* < 0.05), while red dots on the left side indicate miRNAs significantly downregulated (log2 Fold Change < −1.5, FDR *p* < 0.05). Gray dots represent miRNAs that did not meet the significance criteria. The dotted blue lines represent the cut-offs criteria for DE definition.

**Fig. 2:**
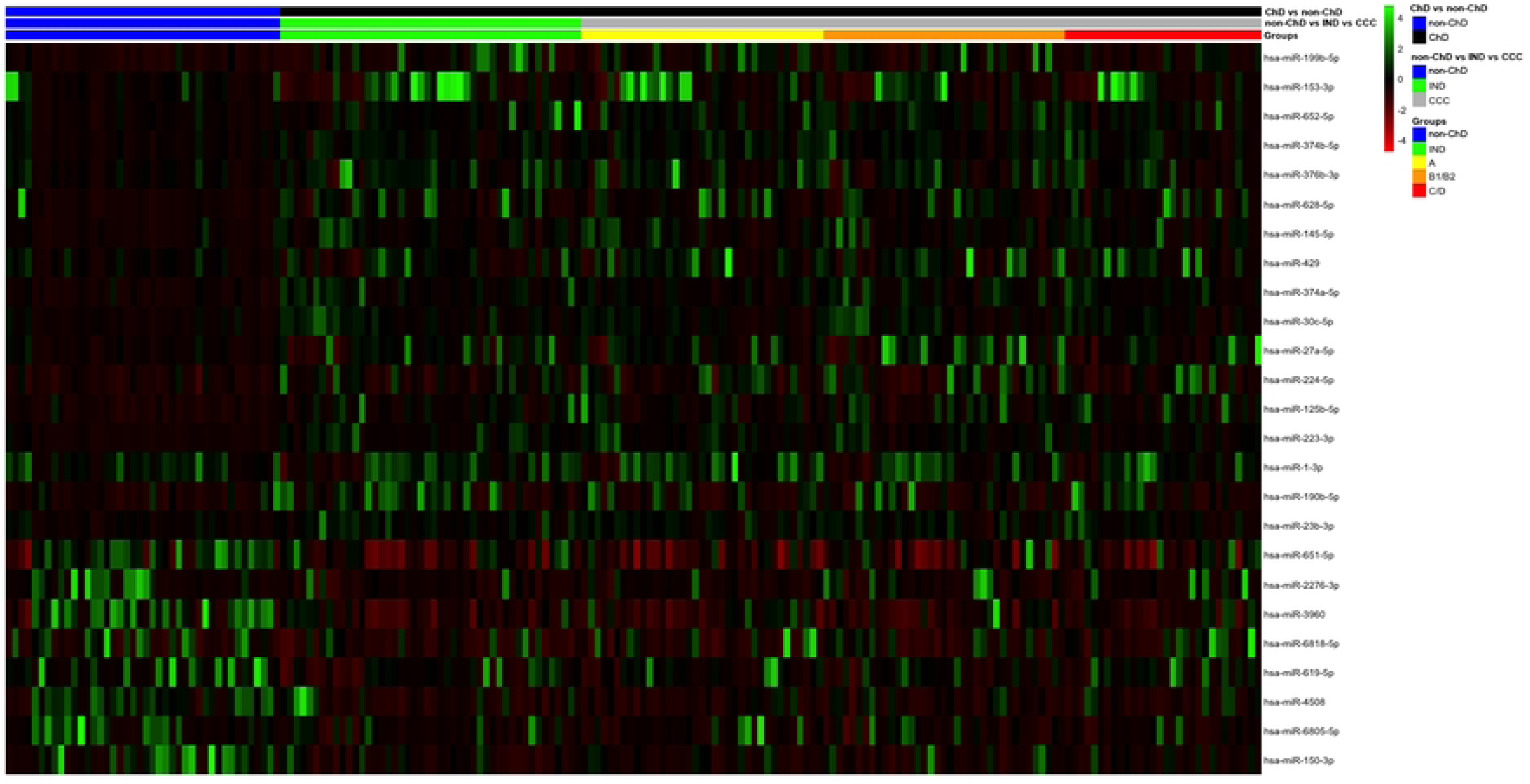
Cluster analysis of differentially expressed miRNAs between Chagas disease (ChD) patients and controls (non-ChD). Heatmap illustrating distinct expression patterns of 17 upregulated and 8 downregulated miRNAs in ChD patients (n=150) compared to non-ChD group (n=42). Each row represents a miRNA, and each column represents a sample. Green indicates higher expression, and red indicates lower expression. non-ChD samples were grouped in blue belt, while ChD group (black belt) and subgroups (Indeterminate, IND, n=46, green belt; CCC, gray belt; A, n=37, yellow belt, B1/B2, orange belt, n=37, and C/D, n=30, red belt) grouped accordingly. The gene expression data was normalized using z-score.

**Fig. 3:**
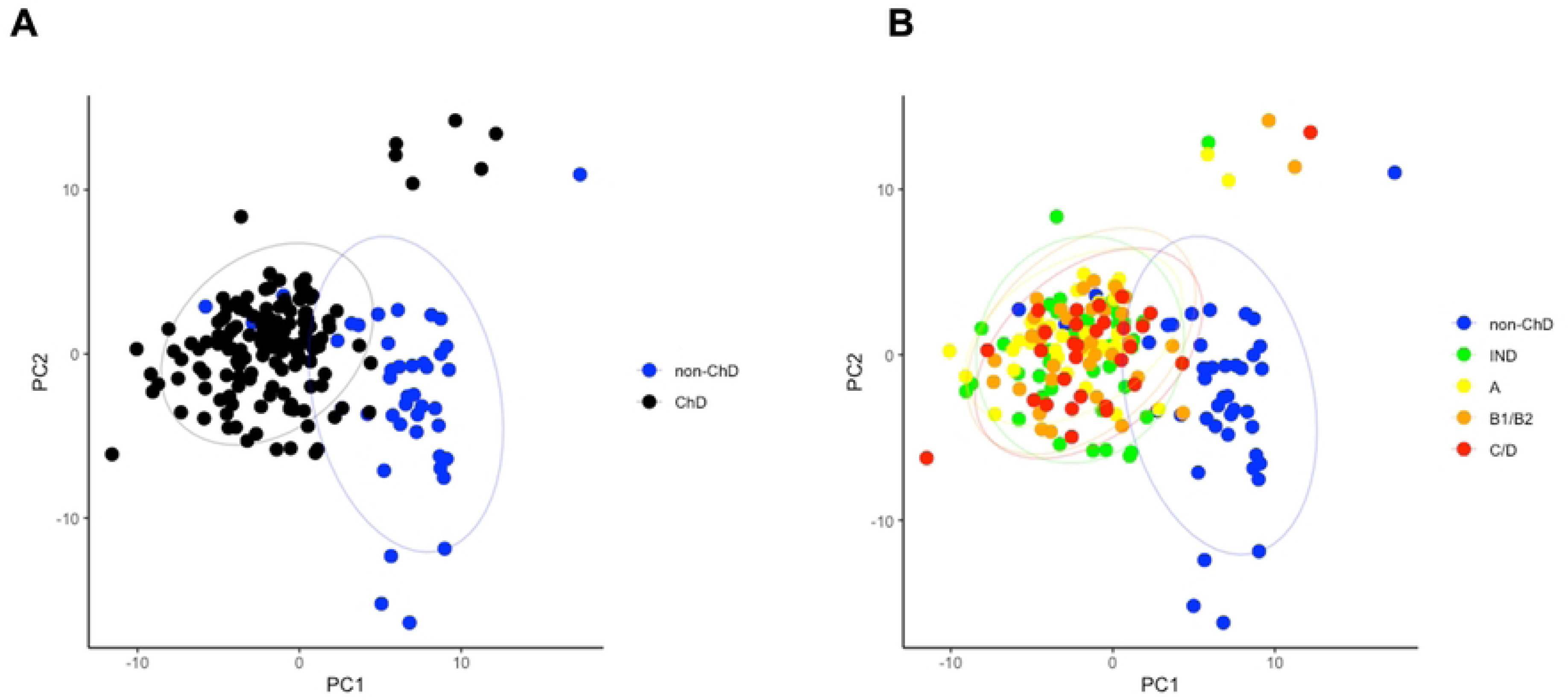
Principal component analysis (PCA) of miRNA expression in Chagas disease (ChD) patients and controls (non-ChD). (A) PCA plot showing a distinguished pattern of differentially expressed miRNAs in ChD patients (black dots, n=150) compared to the non-ChD group (blue dots, n=42). (B) PCA plot showing no differences in miRNA expression between ChD groups (A, n=37, orange, B1/B2, n=37, yellow, C/D, n=30, red, and IND, n=46, green). The colored ellipses represent confidence intervals area, for the respective groups.

As DEMs might be disease biomarkers candidates, we selected 8 (four upregulated and 4 downregulated) of those 40 DEMs in ChD versus non-ChD groups comparison to identify possible differences of expression according to disease form and CCC stage inside ChD subgroups. The eight miRNAs were selected based on highest counts inside DEM criteria, selecting those that could be more feasible to use as biomarkers in diagnostic methods such as qPCR. The analysis of the top 4 upregulated miRNAs with highest counts, miR-143-3p, miR-223-5p, miR-224-5p and miR-454-3p showed significant differences of non-ChD compared to ChD sub-groups. Similar effects were observed in top 4 downregulated miRNAs with highest count, miR-486-5p, miR-629-5p, miR-651-5p and miR-3960, with pronounced significant differences between ChD sub-groups versus non-ChD.

We tested whether DEMs between ChD versus non-ChD could distinguish disease form or CCC stages. However, none of these miRNAs were significantly DEMs in ChD subgroups comparisons in cluster analysis (Fig. 2), In addition, PCA of ChD versus non-ChD DEMs was not able to differentiate ChD subgroups (Fig. 3B). Also, we did not observe statistical differences among ChD subgroups in top upregulated (Fig. 4) or downregulated (Fig. 5) miRNAs expressed in ChD group compared to non-ChD.

**Fig. 4:**
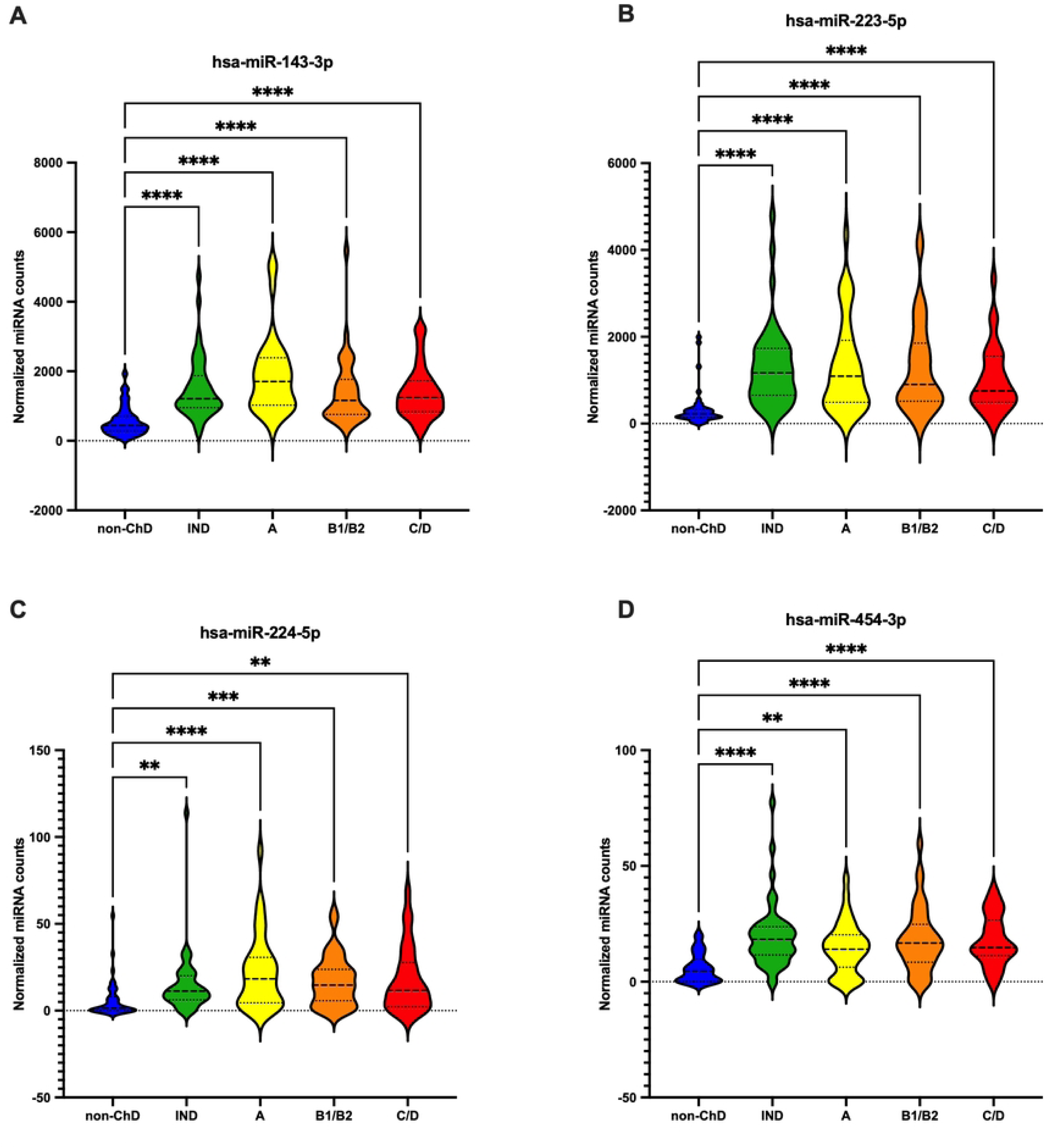
Expression of selected upregulated miRNAs in Chagas disease (ChD) subgroups compared to controls (non-ChD). Violin plots showing the expression levels of miR-143-3p (A), miR-223-5p (B), miR-224-5p (C), and miR-454-3p (D) in the indeterminate form of the disease (IND, n=46), chronic Chagas cardiomyopathy (CCC) stages A (n=37), B1/B2 (n=37), C/D (n=30), and non-ChD group (n=46). The y-axis represents trimmed mean of M-values (TMM) normalized miRNA counts. Statistical significance was determined using the Kruskal-Wallis test corrected with Dunn’s multiple comparisons test. **=*p* < 0.01, ***=*p* < 0.001, ****=*p* < 0.0001.

**Fig. 5:**
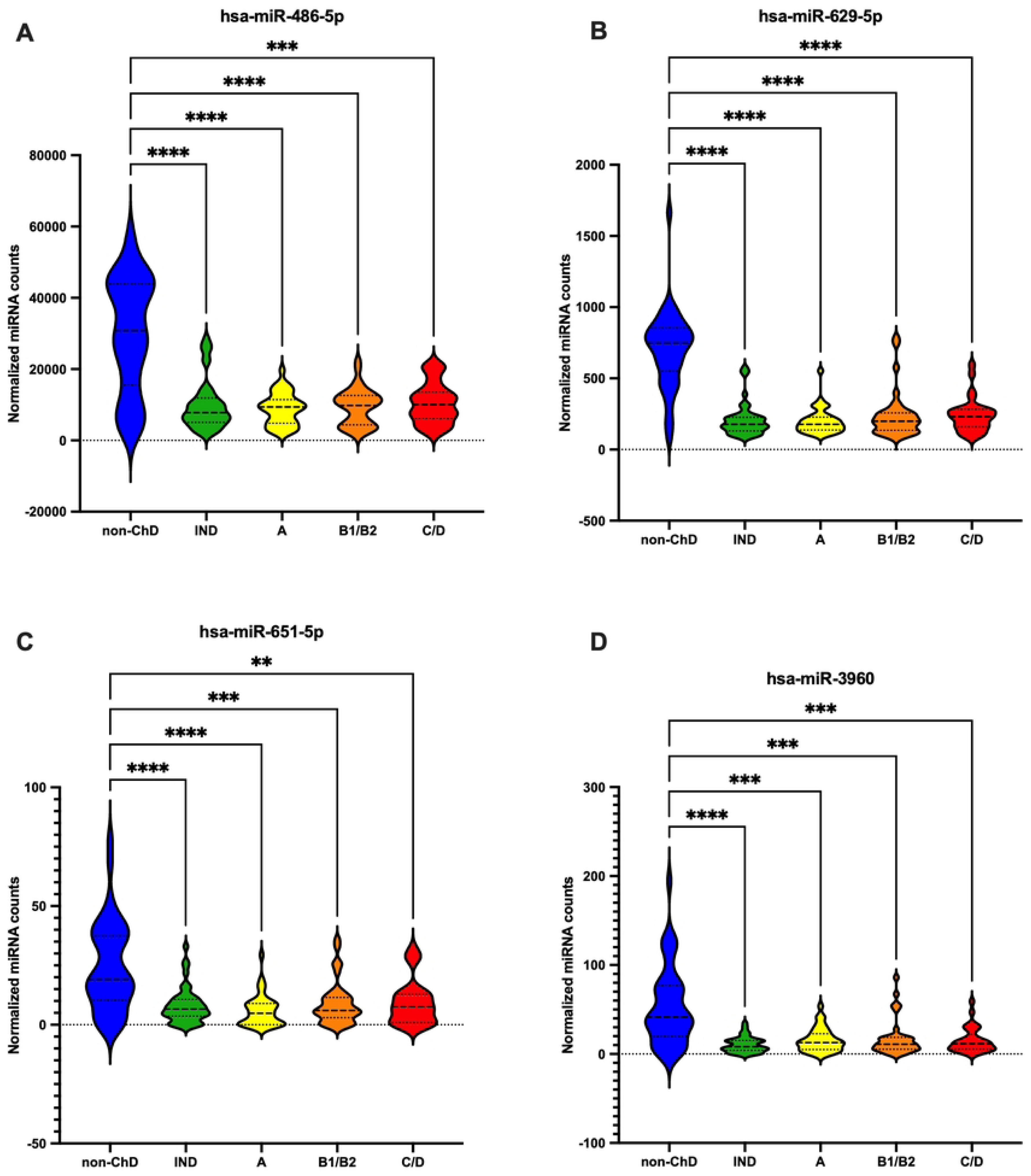
Expression of selected downregulated miRNAs in Chagas disease (ChD) subgroups compared to controls (non-ChD). Violin plots showing the expression levels of miR-486-5p (A), miR-629-5p (B), miR-651-5p (C), and miR-3960 (D) in the indeterminate form of ChD (IND), chronic Chagas cardiomyopathy (CCC) stages A (n=37), B1/B2 (n=37), C/D (n=30), and non-ChD group (n=46). The y-axis represents trimmed mean of M-values (TMM) normalized miRNA counts. Statistical significance was determined using the Kruskal-Wallis test corrected with Dunn’s multiple comparisons test. **=*p* < 0.01, ***=*p* < 0.001, ****=*p* < 0.0001.

### Differential expression analysis between ChD subgroups

We performed differentiation expression analysis between ChD subgroups to identify possible miRNAs able to distinguish ChD forms and CCC stage. We did not observe DEMs between IND and CCC patients. Furthermore, we tested whether miRNAs could be associated with CCC severity, and no differences were observed across CCC groups, as well. Indeed, we did not observe significant differences in miRNAs expression between ChD subgroups. However, analyzing the DEMs using only *p*<0.05 as cutoff, without the FDR correction criteria, it was possible to define some candidates to be associated with CCC stage. The following miRNAs, miR-6734-5p, miR-1285-5p, miR-10527-5p, miR-31-5p, miR-5187-5p, miR-6515-5p were highly expressed in C/D CCC sub-group compared to A CCC sub-group, while miR-30c-2-3p presented lower expression in the same comparison (Fig. 6A). The miRNAs miR-100-5p, miR-642a-3p, and miR-1228-5p were highly expressed, while miR-27b-5p and miR-766-5p were lower expressed in the C/D CCC sub-group compared to B1/B2 CCC sub-group (Fig. 6B). On the other hand, only the miR-10527-5p showed higher expression in the B1/B2 CCC sub-group compared to the A CCC sub-group (Fig. 6C).

**Fig. 6:**
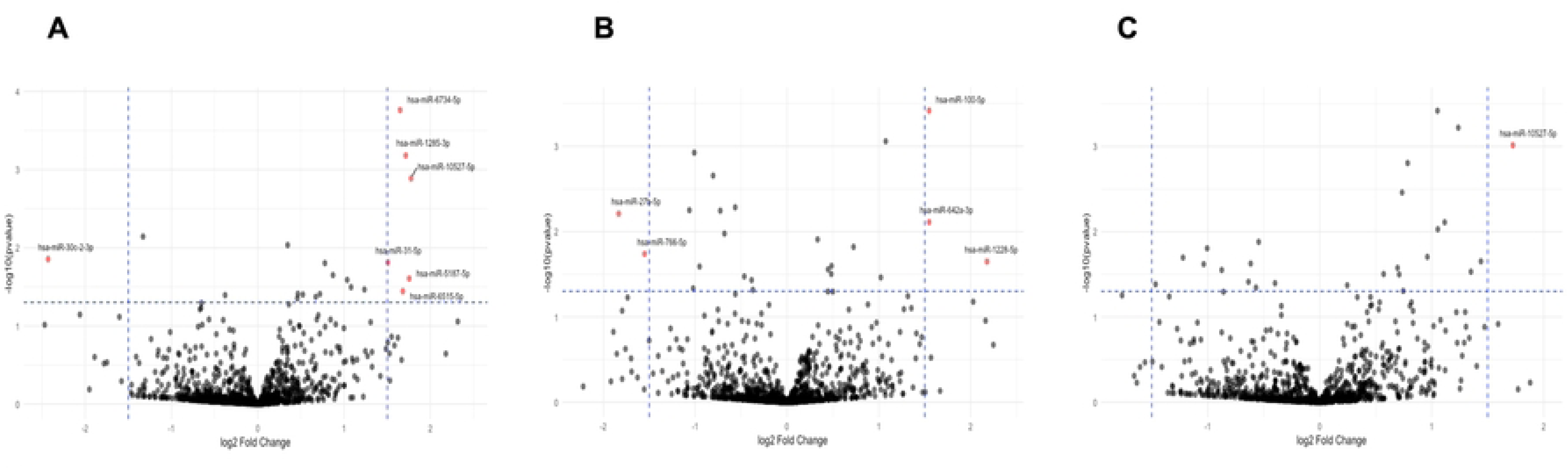
Differentially expressed miRNAs associated with CCC severity. Volcano plots showing differentially expressed miRNAs between CCC groups: (A) C/D-CCC (n=30) versus A-CCC (n=37), (B) C/D-CCC (n=30) versus B1/B2-CCC (n=37), and (C) B1/B2-CCC (n=37) versus A-CCC (n=37). The x-axis represents log2 Fold-change, and the y-axis represents the negative log10 of the p-value. Red dots on the right side of the volcano plot indicate miRNAs significantly upregulated (log2 Fold-change > 1.5, FDR *p* < 0.05), while red dots on the left side indicate miRNAs with significantly downregulated (log2 Fold-change < −1.5, FDR *p* < 0.05). Gray dots represent miRNAs that did not meet the significance criteria. The dotted blue lines represent the cut-offs criteria for DE definition.

To determine the expression levels of these DEMs across CCC stages, we performed an analysis of variance (ANOVA-one way) with counts of these DEMs from CCC sub-groups. The miRNAs miR-6734-5p, miR-1285-3p, miR-10527-5p, and miR-30c-2-3p maintained the significant difference between C/D and A CCC sub-groups (Fig. 7). In addition, the miRNAs miR-100-5p, miR-1228-5p, and miR-766-5p also maintained the significant differences between C/D and B CCC sub-groups (Fig. 8). Interestingly, miR-10527-5p showed a progressive increase in DEM counts from the mild across to the more severe CCC sub-groups (Fig. 6C). Therefore, we performed a Spearman’s correlation analysis to assess the association between miR-10527-5p expression and LVEF, as this is the most important index of CCC severity. We observed a moderate inverse correlation between miR-10527-5p and LVEF percentage (ρ=0.3346, *p*=0.0006) (Fig. 9).

**Fig. 7:**
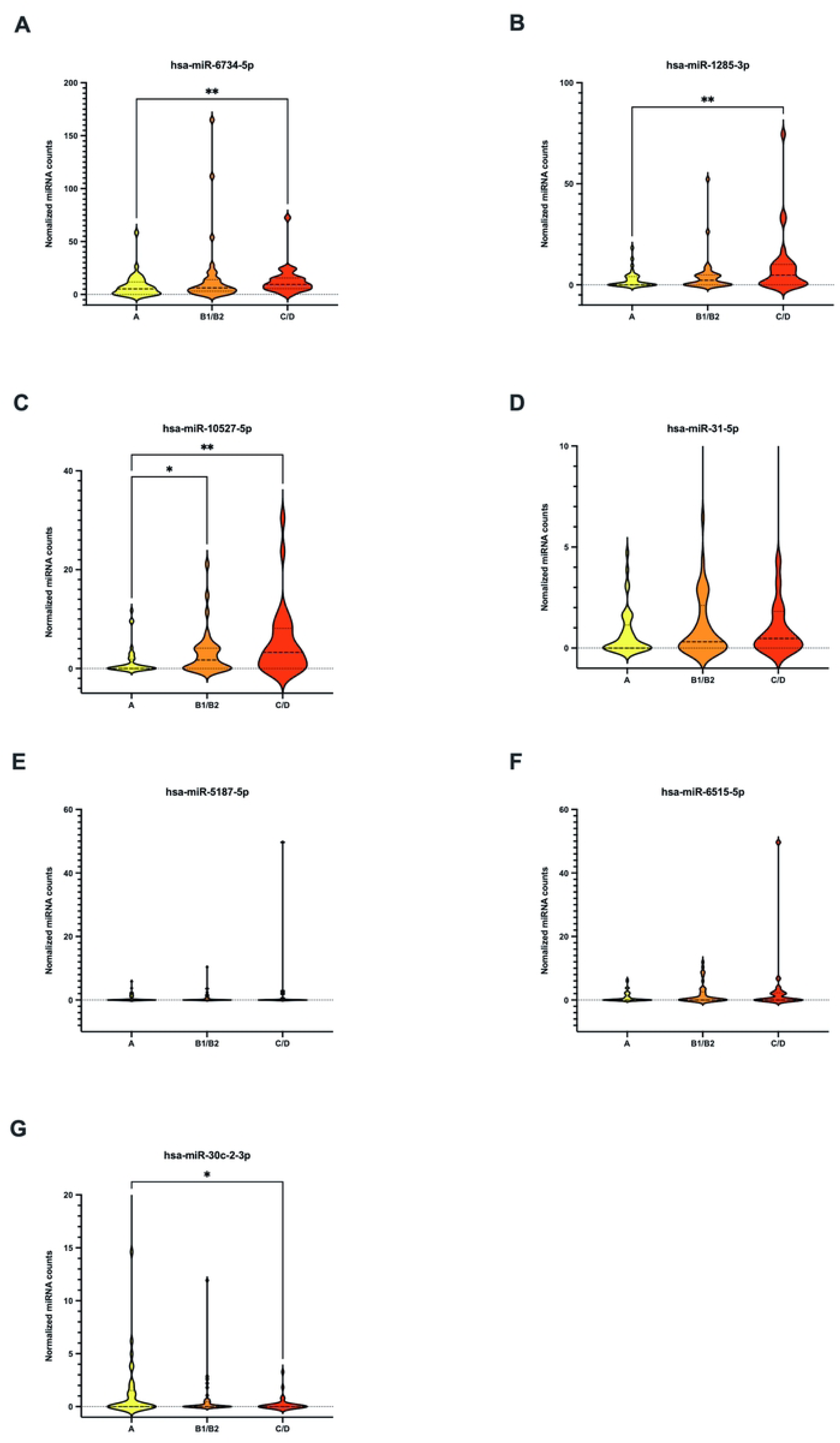
Analysis of differentially expressed miRNAs in C/D versus A-CCC groups. Violin plot showing the expression levels of miR-6734-5p (A), miR-1285-3p (B), miR-10527-5p (C), miR-31-5p (D), miR-5187-5p (E), miR-6515-5p (F), and miR-30c-2-3p (G), in CCC groups A (n=37, yellow), B1/B2 (n=37, orange), C/D (n=30, red). The y-axis represents trimmed mean of M-values (TMM) normalized miRNA counts. Statistical significance was determined using the Kruskal-Wallis test corrected with Dunn’s multiple comparisons test. *=*p* < 0.05, **=*p* < 0.01.

**Fig. 8:**
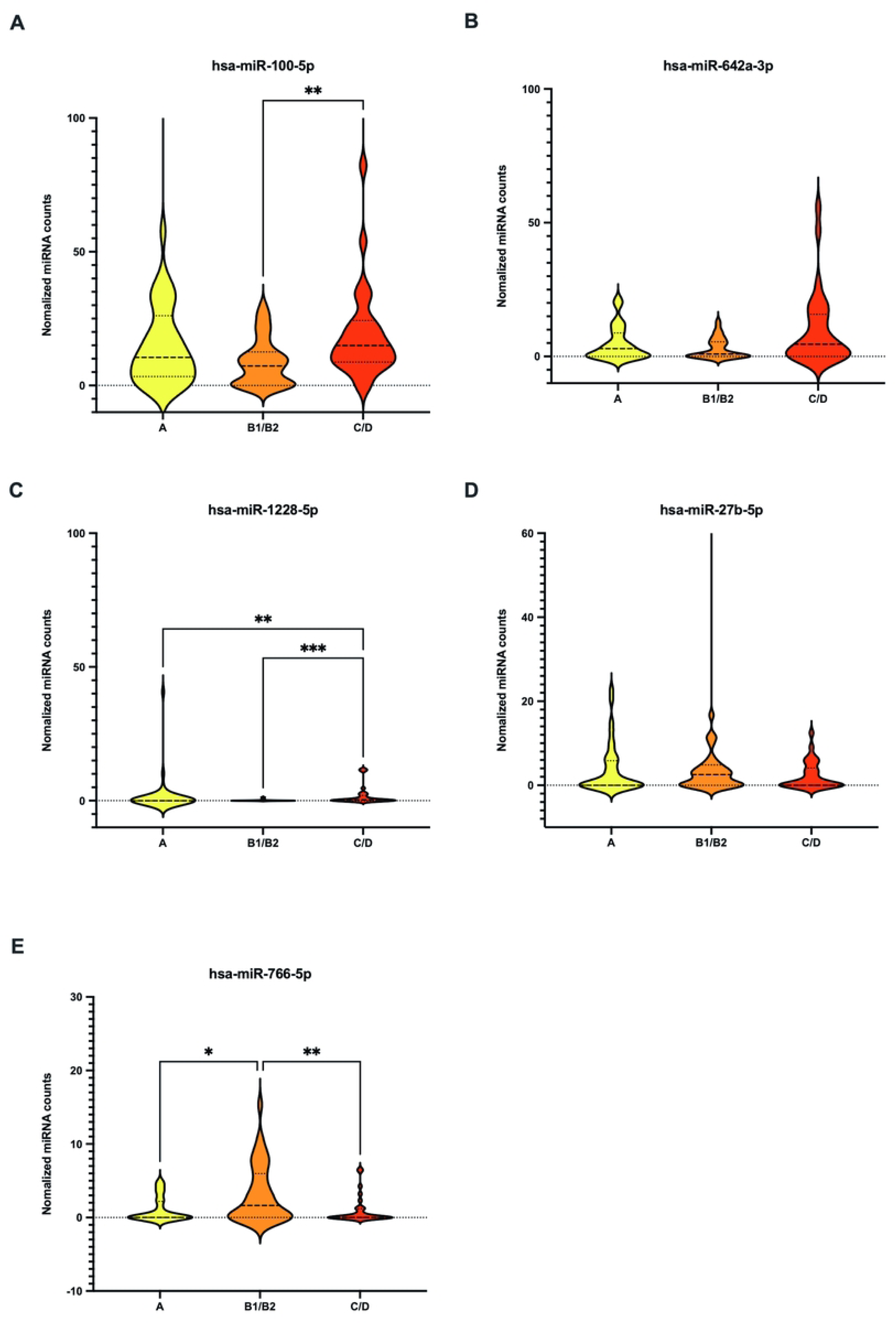
Analysis of differentially expressed miRNAs in C/D versus B1/B2-CCC groups. Violin plot showing the expression levels of miR-100-5p (A), miR-miR-642a-3p (B), miR-1228-5p (C), miR-27b-5p (D), and miR-766-5p, in CCC groups A (n=37, yellow), B1/B2 (n=37, orange), C/D (n=30, red). The y-axis represents trimmed mean of M-values (TMM) normalized miRNA counts. Statistical significance was determined using the Kruskal-Wallis test corrected with Dunn’s multiple comparisons test. *=*p* < 0.05, **=*p* < 0.01, ***=*p* < 0.001.

**Fig. 9:**
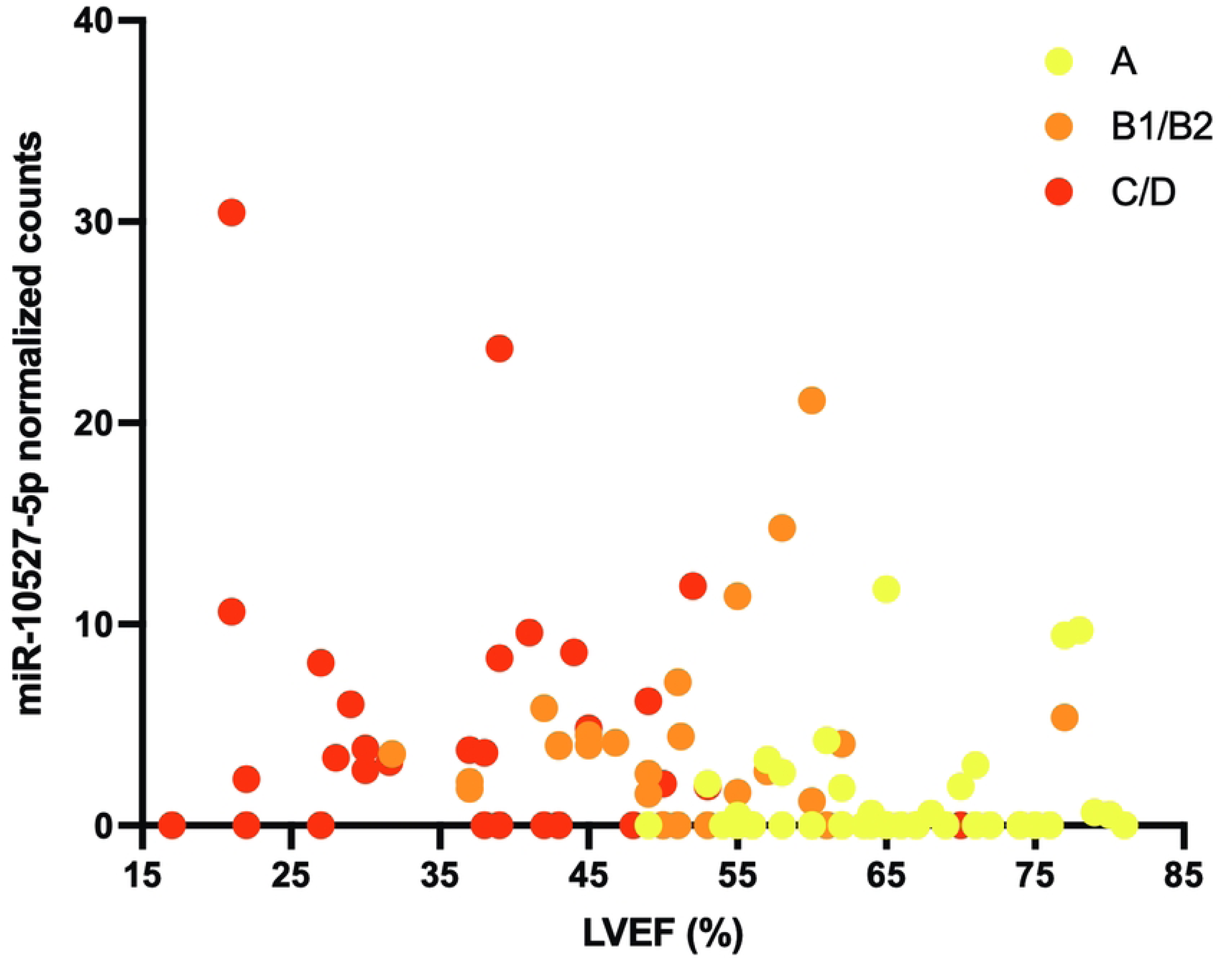
Spearman’s correlation between miR-10527 expression and left ventricular ejection fraction (LVEF) in CCC patients. Dispersion plot showing correlation between miR-10527 TMM normalized counts (y-axis) and LVEF (x-axis) in CCC groups A (n=37, yellow), B1/B2 (n=37, orange), C/D (n=30, red).

### Functional analysis of DEMs in Chagas disease

We performed a functional analysis, by bioinformatics, of the eight miRNAs with functional annotation in the Ingenuity Pathway Analysis (IPA) software from Qiagen (Hilden, Germany), that presented the highest difference of log2 fold-change, and lower FDR p-value in ChD versus non-ChD comparison (Fig. 1), regardless sequencing counts. This analysis was focused on the possible role of these miRNAs regulating immune function and cardiac signaling in ChD, since they presented the highest differences of expression or the highest statistical difference between ChD versus non-ChD groups, independent of their possible use as biomarkers for ChD. The upregulated miRNAs miR-199b-5p and miR-153-3p (highest log2 fold-change value), miR-143-3p and miR-223-3p (lowest FDR p-value), and the downregulated miRNAs miR-150-3p and miR-4508 (highest negative log2 fold-change value), miR-486-5p and miR-3960 (lowest FDR p-value) were selected according to these criteria (Fig 1).

The IPA focusing on the miRNAs interfering pathways related to immune response and cardiovascular diseases and signaling. The IPA was used to explore downstream targets of these selected miRNAs. First, we created a dataset using target scan tool in IPA selecting gene targets for each of those selected miRNAs, filtering for confirmed targets by experimental studies database and targets with highly predicted confidence for miRNA interaction. After target determination, interactions and relationships with biological pathways and diseases related to immune response or cardiovascular signaling were performed.

As miR-199b-5p belongs to the same family of miR199a-5p, with high sequence similarity, we used miR199a-5p as a model for target and biological processes prediction, since miR-199b-5p is not functionally listed in IPA database. The upregulation of miR199a-5p in the ChD group predicted the downregulation in the expression of 51 target genes related to the immune response signaling, including the downregulation of cell proliferation, differentiation, number, and development of lymphocytes (Supplemental Fig 1A). In addition, 21 targets related to cardiovascular signaling, including myocardial infarction and cardiac hypertrophy, were downregulated by miR-199a-5p overexpression in ChD group (Supplemental Fig 1B).

The overexpression of miR-153-3p in ChD patients may be related to downregulation of 33 immune response signaling target genes interacting with decreasing lymphocytes proliferation, differentiation, and cell numbers (Supplemental Fig 2A). Nineteen possible downregulated cardiovascular signaling target genes were observed, promoting cardiac diseases (Supplemental Fig 2B).

The overexpression of miR-143-3p in ChD group predicted downregulation in 48 immune response signaling target genes, influencing the interference in the activation of B and T lymphocytes as well as decreasing in the development of Th1 cells (Supplemental Fig 3A). For cardiovascular signaling, 12 target genes were predicted to be downregulated possibly increasing cardiomyocyte cell death (Supplemental Fig 3B).

Forty genes related to immune response signaling were identified as being downregulated by miR-223-3p overexpression in the ChD group, causing negative effect on lymphocyte numbers, differentiation and expansion, and inflammatory response (Supplemental Fig 4A). Thirteen cardiovascular signaling target genes were identified as potential targets for miR-223-3p downregulation, contributing to dilated cardiomyopathy, acute myocardial infarction (AMI), cardiac fibrosis among other cardiac diseases (Supplemental Fig 4B).

The downregulation of miR-150-3p in ChD patients predicted overexpression in 30 genes related to immune response signaling (Supplemental Fig 5A). The overexpression of these genes is related to increased function, number and activation of lymphocytes, macrophages and plasmacytoid dendritic cells (Supplemental Fig 5B).

The miR-4508 was downregulated in ChD patients and may be related to overexpression of 33 genes related to immune response signaling, promoting differentiation of Th1 cells, activation and increasing numbers of T lymphocytes (Supplemental Fig 6A). Despite the downregulation of the miR-4508 in ChD group predicted overexpression of 10 genes related to cardiovascular signaling, any cardiac disease was predicted (Supplemental Fig 6B).

The downregulation of miR-486-5p in the ChDs group predicted overexpression of 25 genes related to immune response signaling, increasing numbers of leukocytes, and T cell homeostasis (Supplemental Fig 7A). For genes related to cardiovascular signaling, the miR-486-5p downregulation is predicted to upregulate 8 genes that could be strongly related to the development of cardiomyopathy, hypertrophy, and apoptosis of cardiomyocytes (Supplemental Fig 7B).

The miR-3960 downregulated in ChD patients was predicted to overexpress 31 immune response signaling related genes, causing increasing migration and proliferation of lymphocytes, and reducing activation of T reg cells (Supplemental Fig 8A). On the other hand, 15 cardiovascular signaling related genes were predicted to be upregulated in ChDs patients due to miR-3960 downregulation, with mixed effects in the development of cardiac diseases (Supplemental Fig 8B).

We performed similar IPA in the DEMs identified in CCC groups comparison. Only the miRNAs miR-1285-3p, miR-1228-5p and miR-27b showed functional annotation related to immune response signaling and cardiovascular signaling. Forty-three immune response signaling related genes were predicted to be downregulated by miR-1285-3p, miRNA upregulated in C/D-CCC group compared to A-CCC group, influencing negatively in proliferation of T lymphocytes, differentiation of mononuclear leukocytes and in the number of T and Th17 cells (Supplemental Fig 9A). Also, miRNA-1285-3p was predicted to downregulate 22 genes related to cardiovascular signaling, interfering in many cardiac diseases (Supplemental Fig 9B).

The miR-1228-5p was upregulated in C/D-CCC compared to B-CCC group, predicting to downregulate 20 genes related to immune response signaling with possible negative effects on leukocyte migration, macrophage cell movement, and recruitment of myeloid cells (Supplemental Fig 10A). Despite this miRNA was predicted to downregulate 6 genes related to cardiovascular signaling, there is no strong prediction to interfere with cardiac damage events (Supplemental Fig 10B). On the other hand, the downregulation of miR-27-5p in C/D-CCC compared to B-CCC group promotes upregulation of sixteen genes related to immune response signaling that were linked to predict enhanced activation of lymphocytes and T lymphocytes priming, as well as increasing cell viability of leukocytes, apoptosis of dendritic cells and recruitment of lymphocytes and cell cycle progression of T lymphocytes. Interestingly, the miR-27b-5p was predicted to have a negative impact on proliferation of cytotoxic and effector memory T cells (Supplemental Fig 11A). However, we could not identify strong predictions of cardiac signaling events triggered by miR-27b-5p downregulated related genes (Supplemental Fig 11B).

## Discussion

Our study identified a distinct signature of differentially expressed miRNAs in circulating blood of ChD patients compared to non-ChD and within different CCC groups. Specifically, we observed significant upregulation of miR-199b-5p, miR-153-3p, miR-143-3p, and miR-223-3p, and downregulation of miR-150-3p, miR-4508, miR-486-5p, and miR-3960 in ChD versus non-ChD groups. Functional analysis revealed that these miRNAs are involved in pathways related to immune response signaling and cardiovascular signaling, potentially influencing lymphocyte function and the development of cardiac diseases. Notably, upregulated miRNAs in ChD patients are predicted to downregulate genes involved in lymphocyte function and cardiac disease development, while downregulated miRNAs are predicted to have the opposite effect, by upregulating genes related to increased lymphocyte function. In the CCC groups comparison, we identified DE miRNAs like miR-1285-3p, miR-1228-5p and miR-27b that showed functional annotation related to immune response signaling and cardiovascular signaling.

While several studies have mostly explored selected miRNA expression in circulating blood of ChD patients, the application of high-throughput sequencing approaches remains limited. To our knowledge, only one prior study [18] has performed small RNA-Seq to analyze circulating miRNAs in ChD. Villar et al. [18] identified a distinct circulating microRNA signature in chronic ChD. Our study corroborates the involvement of the miR-224-5p but also identifies a novel set of DE miRNAs. These discrepancies could arise from differences in the number of subjects included in the studies (192 in our study versus 66 subjects of Villar et al study), in the study populations (Bolivians versus Brazilians), and data analysis pipelines.

Several studies have highlighted the potential of circulating miRNAs as biomarkers for ChD [16,21]. Prior researches have proposed miR-208a [15] and miR-146a [16] as potential biomarker candidates of indeterminate form of ChD. However, in our study these miRNAs were not able to differentiate IND from CCC patients. Different approaches such as microRNome versus pointed miRNA analysis may explain these different results. Indeed, in the light of our knowledge, our study presented the circulating microRNome analysis of the largest number of patients with ChD to date. Our study was the first to show upregulation of miR-199b-5p, miR153-3p, miR-143-3p, miR-223-3p and downregulation of miR-150-3p, miR-4508, miR-486-5p and miR-3960 in ChD patients compared to non-ChD from a ChD compared to non-ChD from a ChD endemic area.

Although no differences were observed between patients with CCC and those with IND form of ChD, some miRNAs showed altered expression in CCC groups. Duygu et al. [22] showed that miR-199b-5p was involved in cardiac dysfunction and dilation in mouse model of myocardial infarction, corroborating our findings in the functional analysis. In addition, Yang et al. [23] showed increased apoptosis of cardiomyocytes related to the miR-153-3p overexpression. Huang et al. [24] and Scărlătescu et al. [25] proposed the use of miR-143-3p and miR-223-3p, respectively, as biomarkers for sudden cardiac death in acute coronary syndrome and myocardial infarction, corroborating our observations that those miRNAs may play a role in cardiomyocytes cell death and heart diseases. Interestingly, Ribeiro et al. [26] showed that lower levels of miR-223-5p were associated with the worsening of CCC. This seemingly contradictory finding highlights the complexity of miRNA regulation and its context-dependent effects. Contrasting to our results, Hsu et al. [27] and Li et al. [28] showed increased expression of circulating miR-150-3p in patients with cardiac damage, although caused by acute myocardial infarction. However, Li et al. pointed that miR-150-3p was downregulated by medications for heart disease, a common condition in patients with CCC. In addition, Xu et al. [29] observed decreased expression of miR-486-5p in blood of patients with acute myocardial infarction, agreeing with our observations in ChD patients. In contrast with our results, some reports showed increased levels of miR-4508 [30], miR-486-5p [31], and miR-3960 [32] correlating with cardiac abnormalities. Additional studies of these miRNAs in affecting cardiac diseases in the context of ChD deserves more attention. Although no differences were observed between patients with CCC and those with IND form of ChD, some miRNAs showed altered expression in CCC groups. Duygu et al. [22] showed that miR-199b-5p was involved in cardiac dysfunction and dilation in mouse model of myocardial infarction, corroborating our findings in the functional analysis. In addition, Yang et al. [23] showed increased apoptosis of cardiomyocytes related to the miR-153-3p overexpression. Huang et al. [24] and Scărlătescu et al. [25] proposed the use of miR-143-3p and miR-223-3p, respectively, as biomarkers for sudden cardiac death in acute coronary syndrome and myocardial infarction, corroborating our observations that those miRNAs may play a role in cardiomyocytes cell death and heart diseases. Interestingly, Ribeiro et al. [26] showed that lower levels of miR-223-5p were associated with the worsening of CCC. This seemingly contradictory finding highlights the complexity of miRNA regulation and its context-dependent effects. Contrasting to our results, Hsu et al. [27] and Li et al. [28] showed increased expression of circulating miR-150-3p in patients with cardiac damage, although caused by AMI. However, Li et al. pointed that miR-150-3p was downregulated by medications for heart disease, a common condition in patients with CCC. In addition, Xu et al. [29] observed decreased expression of miR-486-5p in blood of patients with AMI, agreeing with our observations in ChD patients. In contrast with our results, some reports showed increased levels of miR-4508 [30], miR-486-5p [31], and miR-3960 [32] correlating with cardiac abnormalities. Additional studies of these miRNAs in affecting cardiac diseases in the context of ChD deserves more attention.

The miRNAs miR-6734-5p, miR-miR-1285-3p, miR-10527-5p, and miR-30c-2-3p were overexpressed in C/D compared to A CCC sub-group. Considering our knowledge, our study was pioneer to show that miR-6734-5p, miR-10527-5p and miR-30c-2-3p were upregulated in severe cardiac disease caused by *T. cruzi* infection. Li et al. [33] have shown the upregulation of miR-1285-3p in plasma and heart of patients during heart failure with strong discrimination from controls, corroborating our results of overexpression of this miRNA under severe cardiomyopathy.

We observed an increased expression of miR-100-5p, miR-1228-5p, and miR-766-5p in C/D compared to B1/B2 CCC group. Zeng et al. [34] showed that overexpression of miR-100-5p induced the expression of cardiac hypertrophy markers in rat model of abdominal aortic coarctation-induced cardiac hypertrophy, by targeting mTOR autophagy activation. In addition, Zhong et al. [30] showed upregulation of circulating miR-1228-5p in patients with acute myocardial infarction. Regarding miR-766-5p, there were no reports attributing this miRNA to cardiac damage or diseases. Taking together, these studies agreed with our attributing the expression of these miRNAs to cardiac damage.

Despite the important contribution to the field, our study has some limitations. First, although this study included the largest number of ChD patient samples for determining miRNAs associated with ChD susceptibility and CCC severity, other studies enrolling Brazilian population from different geographic areas should be addressed to cover all genetic variation that may occur regarding the multiethnic colonization of Brazil. The sample size was still relatively small and derived from a specific Brazilian population, which may limit the generalizability of the findings to broader contexts. Second, the cross-sectional study design restricted our ability to establish the CCC progression outcome. Longitudinal cohort studies may fill this gap by validating these miRNAs as biomarkers of CCC progression. Future studies with larger samples size, more diverse genetic background in the studied population, and longitudinal designs are warranted to validate and expand upon our findings.

In conclusion, our study provides further evidence for the dysregulation of specific miRNAs in ChD. Further research is needed to elucidate the precise roles of these miRNAs in the pathogenesis of ChD and to validate their potential as biomarkers for diagnosis, prognosis, or monitoring disease progression.

## Declarations

### Ethics approval and consent to participate

Written informed consent was obtained from all participants before their participation. This study was approved by the INI/Fiocruz (number 21841519.2.0000.5262) and IOC/Fiocruz (15584119.4.0000.5248) Institutional Review Boards and was conducted according to standards applied by the Brazilian National Committee for Research Ethics and Resolution 466/2012 of the National Health Council, of the Ministry of Health, and to the Declaration of Helsinki, last reviewed in 2024.

### Consent for publication

Not applicable.

### Availability of data and materials

Raw sequencing and processed data are deposited in the NCBI Gene Expression Omnibus (GEO) database (GEO GSE299582).

### Author contributions

EHR, MGBA and RMS conceptualized the study. EHR, MGBA and RMS developed the methodology. FMSA, LFP, AMHM, MTH, LCDS, ARC, FPC, RRF and RMS enrolled participants and collected samples and clinical data. EHR, FMS, FPC and ARSR conducted the experiments. EHR performed the data analysis. MGBA, RMS, JLV and TCAJ acquired funding, with MGBA and RMS supervising the research. The original draft was written by EHR. All authors edited and critically reviewed the manuscript.

## Acknowledgments

We thank Fiocruz RPT01J NGS platform for the support in miRNA library construction and sequencing realized in Rio de Janeiro, Brazil. This work used computational resources provided by the RPT04A Bioinformatics Core Facility at Fiocruz, Rio de Janeiro, Brazil. We thank Renata Malachini Maia for organizing the biorepository and CEPAV-Posse team for their assistance with the non-ChD participants.

---

**Supplemental Fig. 1: Functional analysis of the miR-199a-5p related to immune response and cardiac signaling.** (A) Predictive interaction network of genes related to immune response and miR-199a-5p and possible immune processes and biological pathways. (B) Predictive interaction network of genes related to cardiac signaling and miR-199a-5p and possible influence in cardiac diseases and processes. Symbols, lines colors and formats are descripted in Supplemental Fig. 12.

**Supplemental Fig. 2: Functional analysis of the miR-153-3p related to immune response and cardiac signaling.** (A) Predictive interaction network of genes related to immune response and miR-153-3p and possible immune processes and biological pathways. (B) Predictive interaction network of genes related to cardiac signaling and miR-153-3p and possible influence in cardiac diseases and processes. Symbols, lines colors and formats are descripted in Supplemental Fig. 12.

**Supplemental Fig. 3: Functional analysis of the miR-143-3p related to immune response and cardiac signaling.** (A) Predictive interaction network of genes related to immune response and miR-143-3p and possible immune processes and biological pathways. (B) Predictive interaction network of genes related to cardiac signaling and miR-143-3p and possible influence in cardiac diseases and processes. Symbols, lines colors and formats are descripted in Supplemental Fig. 12.

**Supplemental Fig. 4: Functional analysis of the miR-223-3p related to immune response and cardiac signaling.** (A) Predictive interaction network of genes related to immune response and miR-223-3p and possible immune processes and biological pathways. (B) Predictive interaction network of genes related to cardiac signaling and miR-223-3p and possible influence in cardiac diseases and processes. Symbols, lines colors and formats are descripted in Supplemental Fig. 12.

**Supplemental Fig. 5: Functional analysis of the miR-150-3p related to immune response and cardiac signaling.** (A) Predictive interaction network of genes related to immune response and miR-150-3p and possible immune processes and biological pathways. (B) Predictive interaction network of genes related to cardiac signaling and miR-150-3p and possible influence in cardiac diseases and processes. Symbols, lines colors and formats are descripted in Supplemental Fig. 12.

**Supplemental Fig. 6: Functional analysis of the miR-4508 related to immune response and cardiac signaling.** (A) Predictive interaction network of genes related to immune response and miR-4508 and possible immune processes and biological pathways. (B) Predictive interaction network of genes related to cardiac signaling and miR-4508 and possible influence in cardiac diseases and processes. Symbols, lines colors and formats are descripted in Supplemental Fig. 12.

**Supplemental Fig. 7: Functional analysis of the miR-486-5p related to immune response and cardiac signaling.** (A) Predictive interaction network of genes related to immune response and miR-486-5p and possible immune processes and biological pathways. (B) Predictive interaction network of genes related to cardiac signaling and miR-486-5p and possible influence in cardiac diseases and processes. Symbols, lines colors and formats are descripted in Supplemental Fig. 12.

**Supplemental Fig. 8: Functional analysis of the miR-3960 related to immune response and cardiac signaling.** (A) Predictive interaction network of genes related to immune response and miR-3960 and possible immune processes and biological pathways. (B) Predictive interaction network of genes related to cardiac signaling and miR-3960 and possible influence in cardiac diseases and processes. Symbols, lines colors and formats are descripted in Supplemental Fig. 12.

**Supplemental Fig. 9: Functional analysis of the miR-1285-3p related to immune response and cardiac signaling.** (A) Predictive interaction network of genes related to immune response and miR-1285-3p and possible immune processes and biological pathways. (B) Predictive interaction network of genes related to cardiac signaling and miR-1285-3p and possible influence in cardiac diseases and processes. Symbols, lines colors and formats are descripted in Supplemental Fig. 12.

**Supplemental Fig. 10: Functional analysis of the miR-1228-5p related to immune response and cardiac signaling.** (A) Predictive interaction network of genes related to immune response and miR-1228-5p and possible immune processes and biological pathways. (B) Predictive interaction network of genes related to cardiac signaling and miR-1228-5p and possible influence in cardiac diseases and processes. Symbols, lines colors and formats are descripted in Supplemental Fig. 12.

**Supplemental Fig. 11: Functional analysis of the miR-27-5p related to immune response and cardiac signaling.** (A) Predictive interaction network of genes related to immune response and miR-27-5p and possible immune processes and biological pathways. (B) Predictive interaction network of genes related to cardiac signaling and miR-27-5p and possible influence in cardiac diseases and processes. Symbols, lines colors and formats are descripted in Supplemental Fig. 12.

**Supplemental Fig. 12: Legends for functional analysis figures.** (A) Symbols to identify molecules functions and biological processes. (B) Relationship nodes and interconnecting lines meaning. (C) and (D) Colors identifying the regulation state of the molecule.

